# Integrated behavioural, morphological, and reproductive responses reveal a trade-off during diapause in *Culex pipiens*

**DOI:** 10.64898/2026.06.12.731944

**Authors:** Rosheen S. Mthawanji, Amirah Binti Rashid, Jola Tanianis-Hughes, Krishanthi S. Subramaniam, Marcus S C. Blagrove

## Abstract

Diapause is a critical adaptive strategy that enables temperate mosquito species to survive adverse environmental conditions and maintain population persistence across seasons. In *Culex pipiens*, diapause plays a key role in overwintering and influences the seasonal dynamics of arbovirus transmission. However, diapause expression is often assessed using single traits, limiting our understanding of its integrated physiological basis and variation among populations.

In this study, we investigated the behavioural, morphological, and reproductive signatures of diapause across three laboratory strains of *Culex pipiens* (Mogden, Pirbright, and Pirbright Hybrid) reared under diapause-inducing (10 °C), cold (14 °C), and control (26–27 °C) conditions. We quantified blood-feeding behaviour, wing size as a proxy for somatic growth, and spermatheca size as an indicator of reproductive development.

Diapause-inducing conditions resulted in a coordinated phenotype characterised by strong suppression of blood-feeding, increased somatic size, and marked inhibition of reproductive development. Mosquitoes reared at 10 °C exhibited near-complete feeding inhibition and significantly reduced spermatheca size, consistent with reproductive arrest, while those reared at 14 °C showed intermediate phenotypes. In contrast, control mosquitoes displayed active feeding and fully developed reproductive structures. Wing size increased progressively with decreasing temperature, with the largest individuals observed under diapause-inducing conditions.

When analysed together, wing size and spermatheca development exhibited opposing responses across temperature treatments, revealing a strong negative association and indicating a trade-off between somatic growth and reproductive investment. This integrated response supports the interpretation of diapause as a coordinated life-history strategy involving resource reallocation towards survival.

Additionally, diapause expression varied among strains, with the Mogden strain showing reduced sensitivity compared with Pirbright and hybrid populations, highlighting the role of genetic background in diapause plasticity.

These findings demonstrate that diapause in *Culex pipiens* is a multi-trait, plastic phenotype with important implications for overwintering success and the seasonal dynamics of arbovirus transmission in temperate regions.

## Introduction

Diapause is a key adaptive strategy that enables temperate insect species to survive adverse environmental conditions and persist across seasonal cycles (Denlinger, 2002; Denlinger & Armbruster, 2014). In mosquitoes, diapause is particularly important for overwintering and has significant implications for the seasonal dynamics of arbovirus transmission. In *Culex pipiens*, a primary vector of West Nile virus and Usutu virus, diapause allows populations to persist through winter and reinitiate transmission in spring (Turell, 2012; Farajollahi et al., 2011). Understanding how diapause is expressed is therefore critical for predicting mosquito population dynamics and disease risk, particularly under changing climate conditions.

In *Culex pipiens*, diapause is induced by a combination of low temperature and short photoperiod and is characterised by behavioural and physiological changes, including suppression of blood-feeding, reproductive arrest, altered metabolism, and increased lipid storage (Robich & Denlinger, 2005; Denlinger & Armbruster, 2014). These traits collectively enable females to conserve energy and enhance survival during unfavourable conditions.

While these diapause-associated responses have been well documented, they are often examined in isolation. Despite extensive study, diapause is still predominantly assessed using single traits, limiting our ability to understand it as an integrated life-history strategy shaped by resource allocation trade-offs.

Current understanding of diapause is limited not only by focusing on single-trait assessments, but also by insufficient consideration of how genetic background shapes strain-specific expression. This is important given evidence that genetic background can modulate responsiveness to diapause-inducing cues (Sauer et al., 2022; Nelms et al., 2013). Variation in diapause plasticity among strains retains significant ecological consequences through influencing overwintering survival, the timing of emergence, and ultimately, the seasonal dynamics of arbovirus transmission.

In this study, we investigated the behavioural, morphological, and reproductive signatures of diapause in three laboratory strains of *Culex pipiens* under controlled temperature and photoperiod conditions. Specifically, we quantified blood-feeding behaviour, wing size as a proxy for somatic growth, and spermatheca size as an indicator of reproductive development. By integrating these traits, we aimed to determine whether diapause represents a coordinated life-history strategy characterised by trade-offs between somatic growth and reproduction, and to assess how its expression varies across strains.

## Methods

### Mosquito Strains and Rearing Conditions

Three strains of *Culex pipiens* were used throughout the experiments, representing two biotypes: *Culex pipiens pipiens* and *Culex pipiens molestus.* Biotypes were provisionally classified based on their ecological characteristics, with *Cx. p. pipiens* associated with above-ground (epigean) habitats and *Cx. p. molestus* associated with underground (hypogean) environments (Mogden, UK).

The *Cx. p. pipiens* strain was obtained from the Pirbright Institute (UK). In addition, a hybrid strain derived from the Pirbright colony was included in the study (Manley et al., 2015).

All colonies were maintained under standard insectary conditions (26–27 °C, 70–80% room humidity (RH), 12:12 light: dark) prior to the experiment. Eggs were hatched in dechlorinated tap water, and first instar (L1) larvae were transferred to experimental temperature treatments within 12 hours of hatching.

To evaluate diapause-associated physiological responses, for each treatment, 200 larvae per strain were reared from first instar (L1) to pupation under one of three temperature regimes and photoperiod (Figure 1):

1. Diapause-inducing temperature: 10 °C, 8:16h light: dark cycle
2. Cold temperature: 14 °C, 16:8h light: dark cycle
3. Control temperature: 26–27 °C, 12:12h light: dark cycle

**Figure 1.**
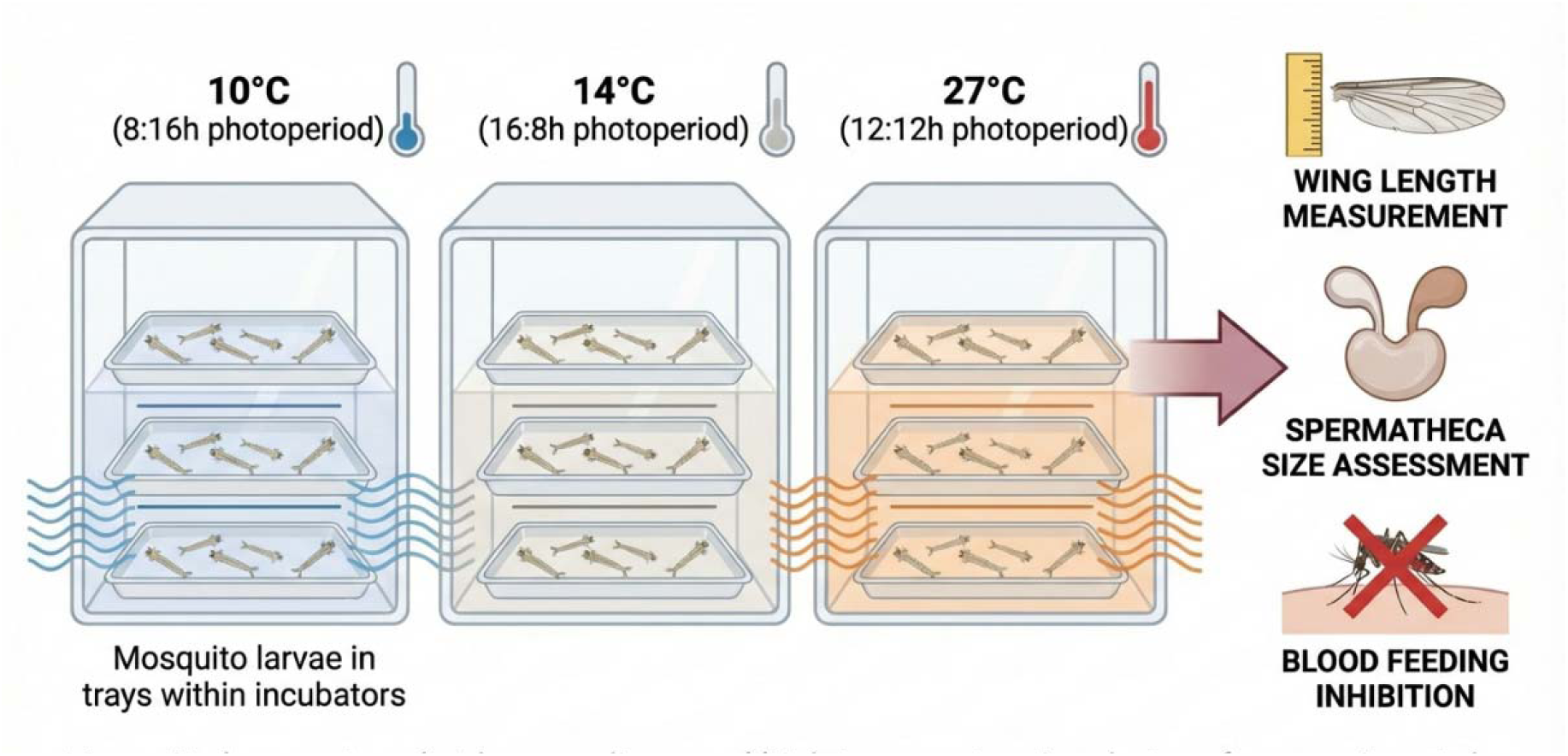
Experimental design and environmental conditions used to induce diapause in *Culex* pipiens. Schematic overview of temperature and photoperiod treatments applied from the first instar larval stage (L1) through to adult emergence. Mosquitoes were reared under three environmental regimes: diapause-inducing conditions (10 °C, 8:16 light: dark), cold conditions (14 °C, 16:8 light: dark), and control insectary conditions (26–27 °C, 12:12 light: dark). Following emergence, adult females were assessed for behavioural (blood-feeding), morphological (wing length), and reproductive (spermatheca size) traits. This design enables comparison of integrated diapause-associated responses across environmental conditions.

**Figure 2.**
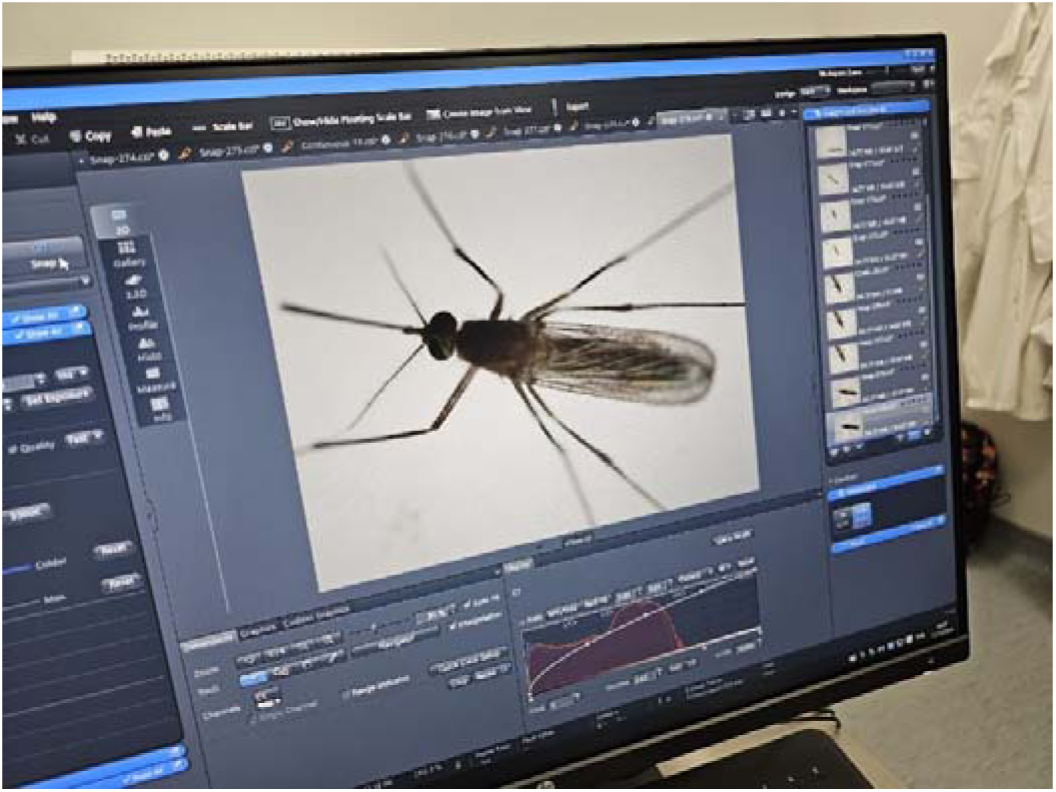
Measurement of wing length as a proxy for somatic size in *Culex* pipiens. Representative image showing the method used to measure wing length under a stereomicroscope (Zeiss Axio Zoom V16) at 40× magnification. Wing length (µm) was measured from the alular notch to the apical tip, excluding fringe scales, using calibrated digital imaging software. Wing size was used as a proxy for overall somatic growth, allowing comparison of developmental responses across temperature treatments.

Photoperiods were selected to reflect established diapause-inducing and non-inducing regimes for *Cx. pipiens*, in which short day length synergises with low temperature to induce adult diapause

### Sample sizes

Sample sizes were determined based on previous studies assessing diapause-associated traits in *Culex pipiens* and other mosquito species, where similar numbers have been shown to provide sufficient statistical power to detect biologically meaningful differences (Robich & Denlinger, 2005; Field et al., 2022). For blood-feeding assays, each treatment consisted of 35 females per strain across 4 independent replicates.

### Blood-Feeding Inhibition Assay

Blood-feeding inhibition assays were conducted using adult female mosquitoes aged 7–10 days post-emergence, corresponding to the typical host-seeking period. Females were maintained in mixed-sex cages prior to the assay and were therefore either mated or unmated.

To standardise feeding motivation, mosquitoes were deprived of sucrose for 24 hours prior to blood feeding. Following starvation, females were offered defibrinated horse blood using a Hemotek membrane feeding system (Graumans *et al*., 2023).

Feeding assays were conducted for 45 minutes under low light. Feeding success was assessed by visually identifying fully engorged females after the feeding period. The proportion of blood-fed females was calculated for each experimental group and pooled across biological replicates.

### Wing Morphometrics

To assess the effects of temperature on somatic growth, wing length was used as a proxy for body size (Field *et al*., 2022). The left wing was removed from each female mosquito, mounted individually on a microscope slide. Wing measurements were obtained using a Zeiss Axio Zoom V16 microscope equipped with calibrated digital image analysis software. Wing length (µm) was measured from the alular notch to the apical tip, excluding fringe scales (Figure 3).

**Figure 3.**
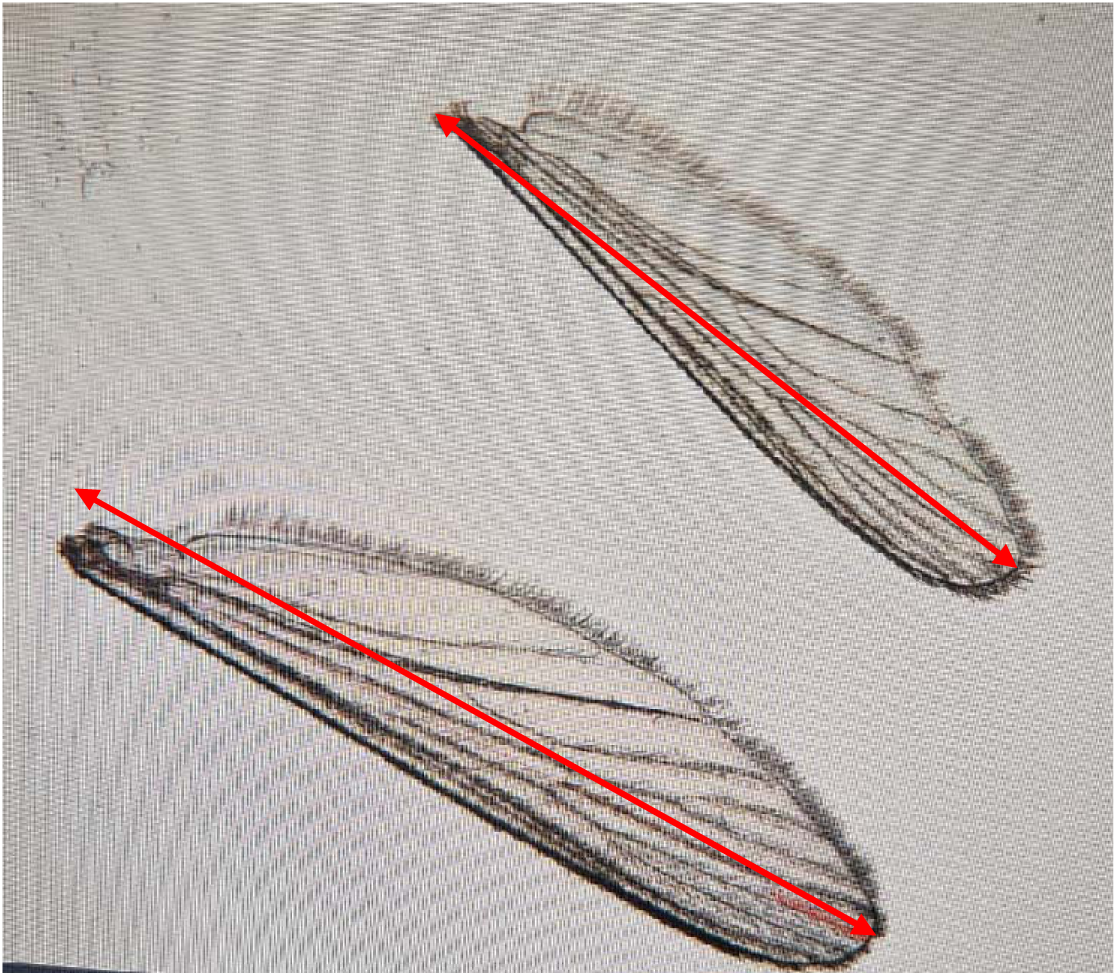
Comparative wing size of *Culex* mosquitoes exposed to different temperature conditions. The larger wing (bottom left) is from an individual exposed to extreme low-temperature diapause conditions, while the smaller wing (top) is from one maintained at normal insectary temperatures. The image highlights the influence of temperature on morphological development

### Spermatheca Dissections and Measurements

To quantify reproductive development, all three spermathecae were dissected from adult females. Spermathecae were removed in cold PBS, transferred to slides, photographed immediately to minimise collapse, and measured using the Zeiss microscope ruler (Figure 4). For each female, the diameter of each spermatheca was measured, and a mean spermatheca size (µm) was calculated per mosquito. Damaged or collapsed spermathecae were excluded. Spermatheca measurements were conducted for all strains.

**Figure 4.**
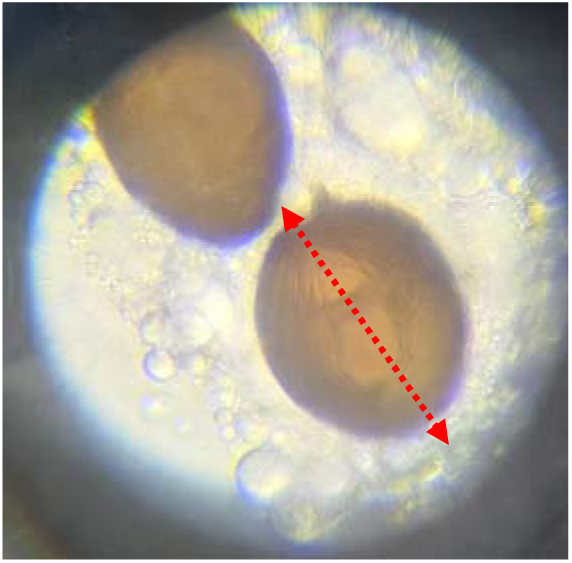
Dissection and measurement of spermathecae as an indicator of reproductive development in *Culex* pipiens. Representative image of dissected spermathecae from an adult female mosquito following exposure to low-temperature conditions. Spermathecae were removed in cold PBS, mounted on slides, and imaged immediately to prevent structural collapse. The diameter (µm) of each spermatheca was measured using calibrated microscopy, and mean spermatheca size per individual was used as a proxy for reproductive development. Reduced spermatheca size is indicative of reproductive arrest associated with diapause.

### Data Analysis

All statistical analyses were performed in R (version 4.3). Data were assessed for normality and homogeneity of variance prior to analysis.

Blood-feeding inhibition was analysed using a two-way ANOVA with temperature treatment (10 °C, 14 °C, 26–27 °C) and adult age (7 or 30 days) as fixed factors. Wing length was analysed using a two-way ANOVA with temperature as the fixed factor while spermatheca size was analysed using a one-way ANOVA.

## Results

### Blood-feeding behaviour differed markedly across temperature treatments

Females reared under diapause-inducing conditions (10 °C) exhibited strong suppression of feeding at both 7 and 30 days post-emergence (Figure 5). At 7 days, only a small proportion of diapause-reared females fed, whereas more than 70% of females in both the cold (14 °C) and control (26–27 °C) treatments successfully took a blood meal. By 30 days, blood-feeding in diapause-reared females was almost completely absent, while feeding rates in cold and control groups remained moderate.

**Figure 5.**
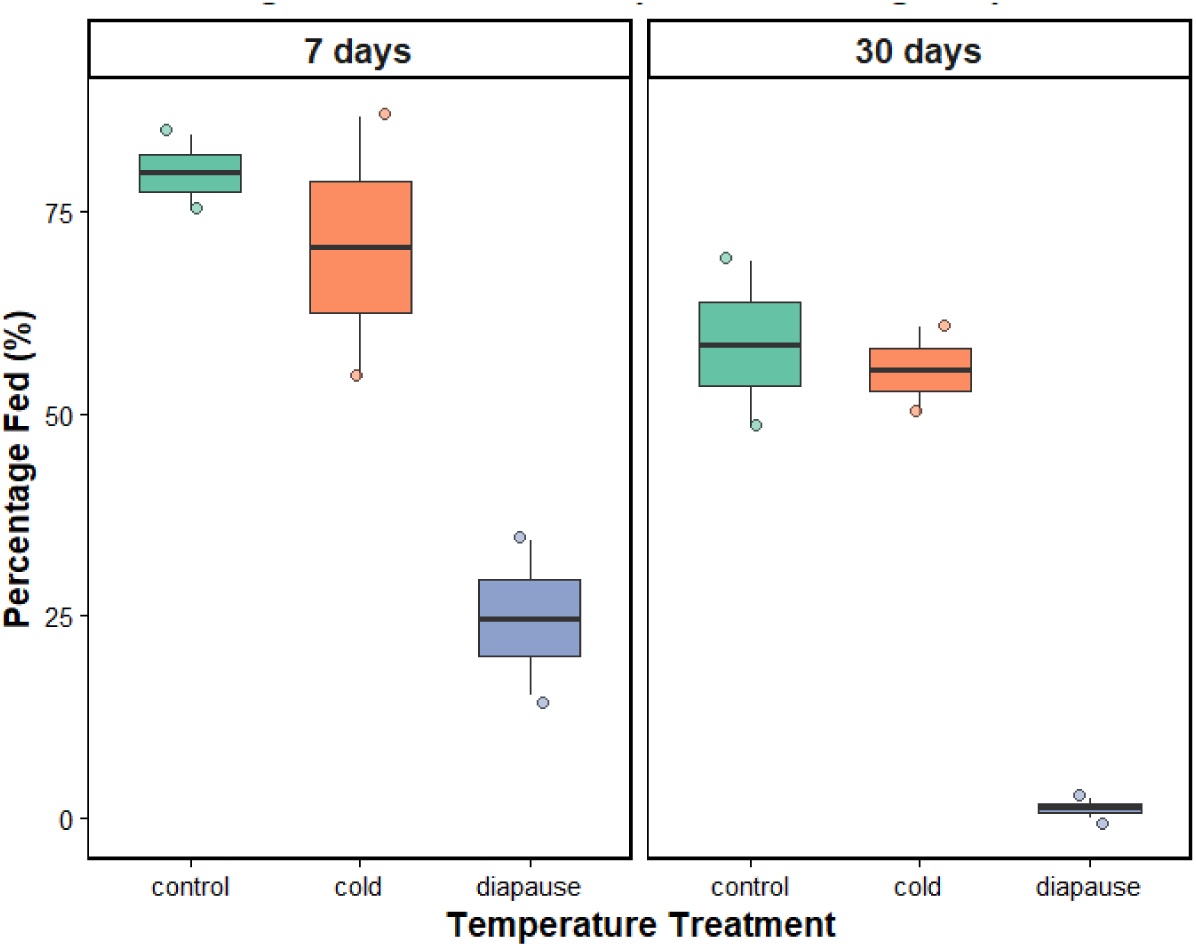
Feeding inhibition under diapause-inducing temperatures. Percentage of females that successfully blood-fed at 7- and 30-days post-emergence when reared at diapause (10 °C), cold (14 °C), or control (26–27 °C) temperatures. Boxplots show the distribution of feeding success, and points represent individual replicates. Feeding was strongly suppressed at 10 °C across both ages, consistent with diapause-associated behavioural arrest

Two-way ANOVA revealed significant effects of temperature treatment (F□,□ = 22.25, p = 0.0017) and age (F□,□ = 7.01, p = 0.038) on feeding success, with no significant interaction between treatment and age (F□,□ = 0.11, p = 0.90). Post hoc comparisons confirmed that feeding rates in the diapause group were significantly lower than in both cold and control treatments (p < 0.01), while no significant difference was observed between cold and control groups.

Wing length varied significantly across temperature treatments and mosquito strains (Figure 6 and 7). Across all strains, mosquitoes reared at 10 °C exhibited the largest wings, those at 14°C showed intermediate sizes, and those at 26–27 °C had the smallest.

**Figure 6.**
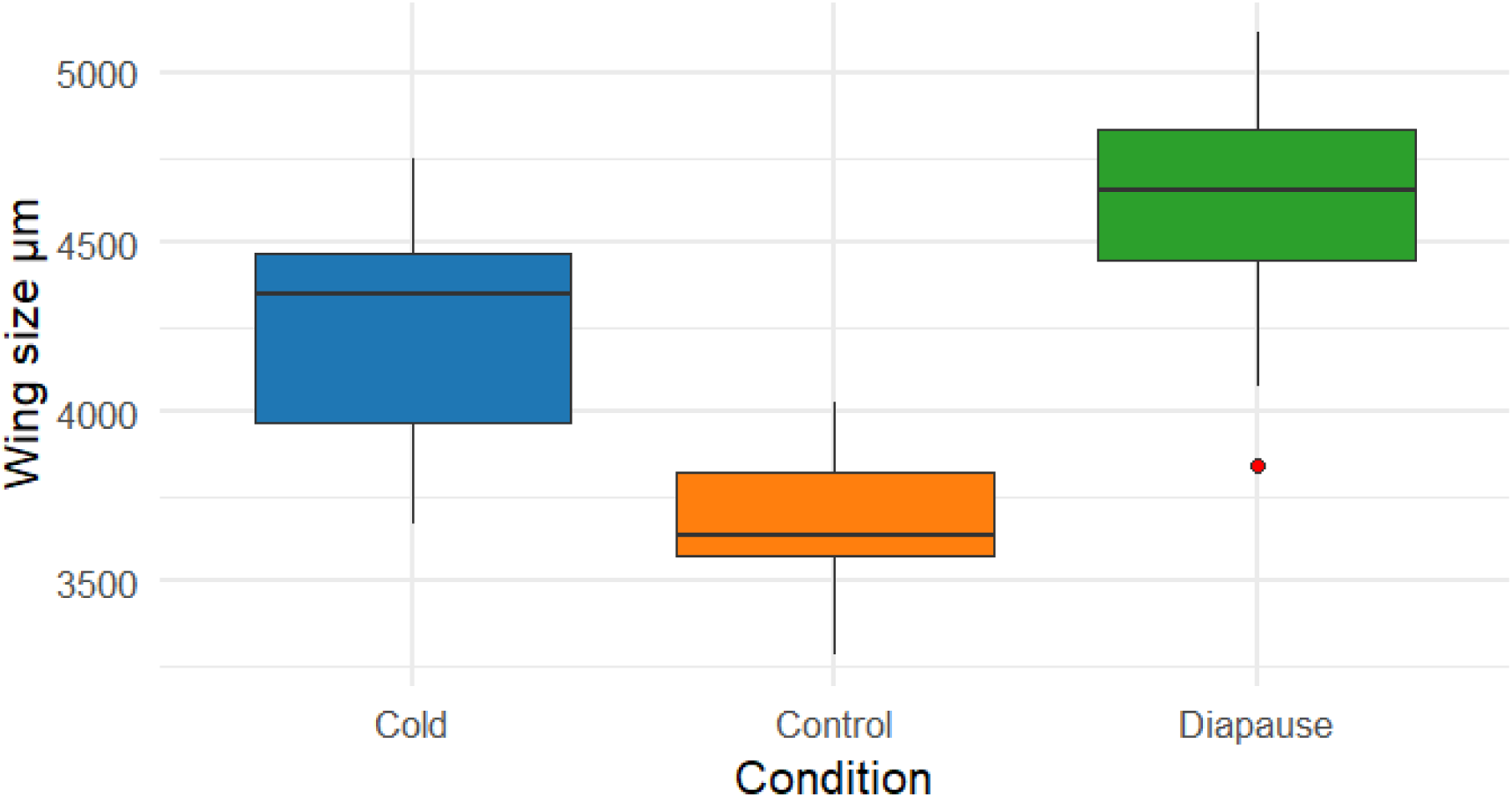
Effect of temperature on wing size in *Culex pipiens*. Wing length (µm) of adult females across diapause (10 °C), cold (14 °C), and control (26–27 °C) conditions, across all strains. Boxplots show the distribution of wing size, with individual points representing mosquitoes. Wing size increased with decreasing temperature, with the largest individuals observed under diapause-inducing conditions.

**Figure 7.**
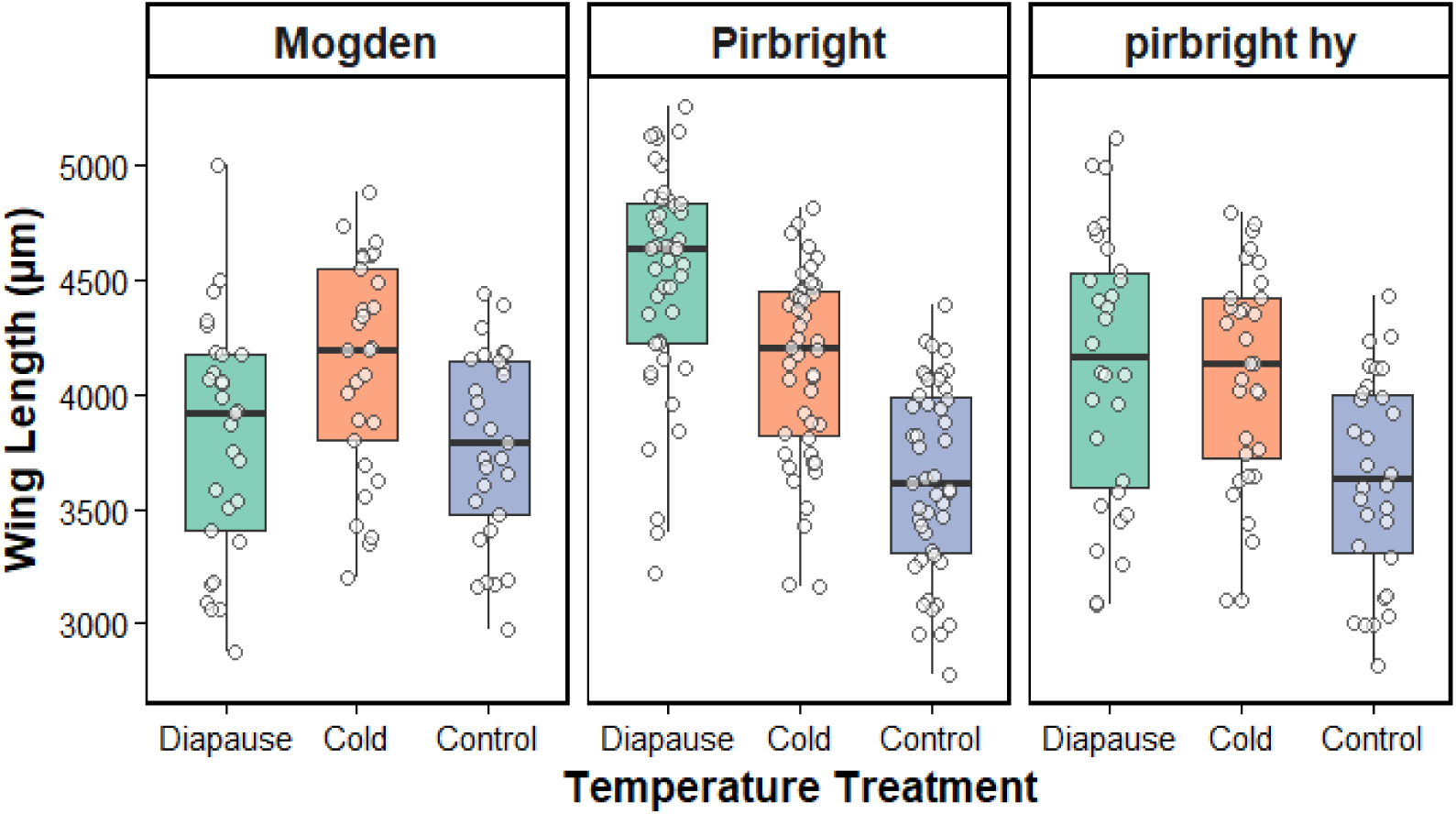
Effect of temperature on wing size across *Culex pipiens* strains. Wing length (µm) of adult females reared at diapause (10 °C), cold (14 °C), or control (26–27 °C) temperatures. Boxplots show somatic growth differences across treatments for the m*olestus,* Pirbright, and Pirbright Hybrid strains. Individual points represent single mosquitoes. Wing size increased with a decline in temperature in Pirbright and Hybrid strains, while the *molestus* strain showed attenuated temperature sensitivity.

Two-way ANOVA revealed significant effects of temperature (F□,□□□ = 40.61, p < 0.001), strain (F□,□□□ = 5.46, p = 0.0047), and a significant treatment × strain interaction (F□,□□□ = 8.77, p < 0.001), indicating strain-specific differences in thermal plasticity.

Post hoc comparisons showed that *Culex pipiens* and *pipiens hybrid* had significantly larger wings than Mogden *molestu*s (mean difference =190 µm, p = 0.008), while the hybrid strain had smaller wings than Pirbright (p = 0.045) and did not differ significantly from the *molestus*.

Within-strain (Figure 8) analyses indicated that temperature effects were most pronounced in Pirbright and hybrid strains, which exhibited clear increases in wing size when temperature was under lower temperature conditions, with the largest wings observed under diapause-inducing conditions. In contrast, the *molestus* showed relatively limited variation across treatments, suggesting reduced sensitivity to temperature.

**Figure 8.**
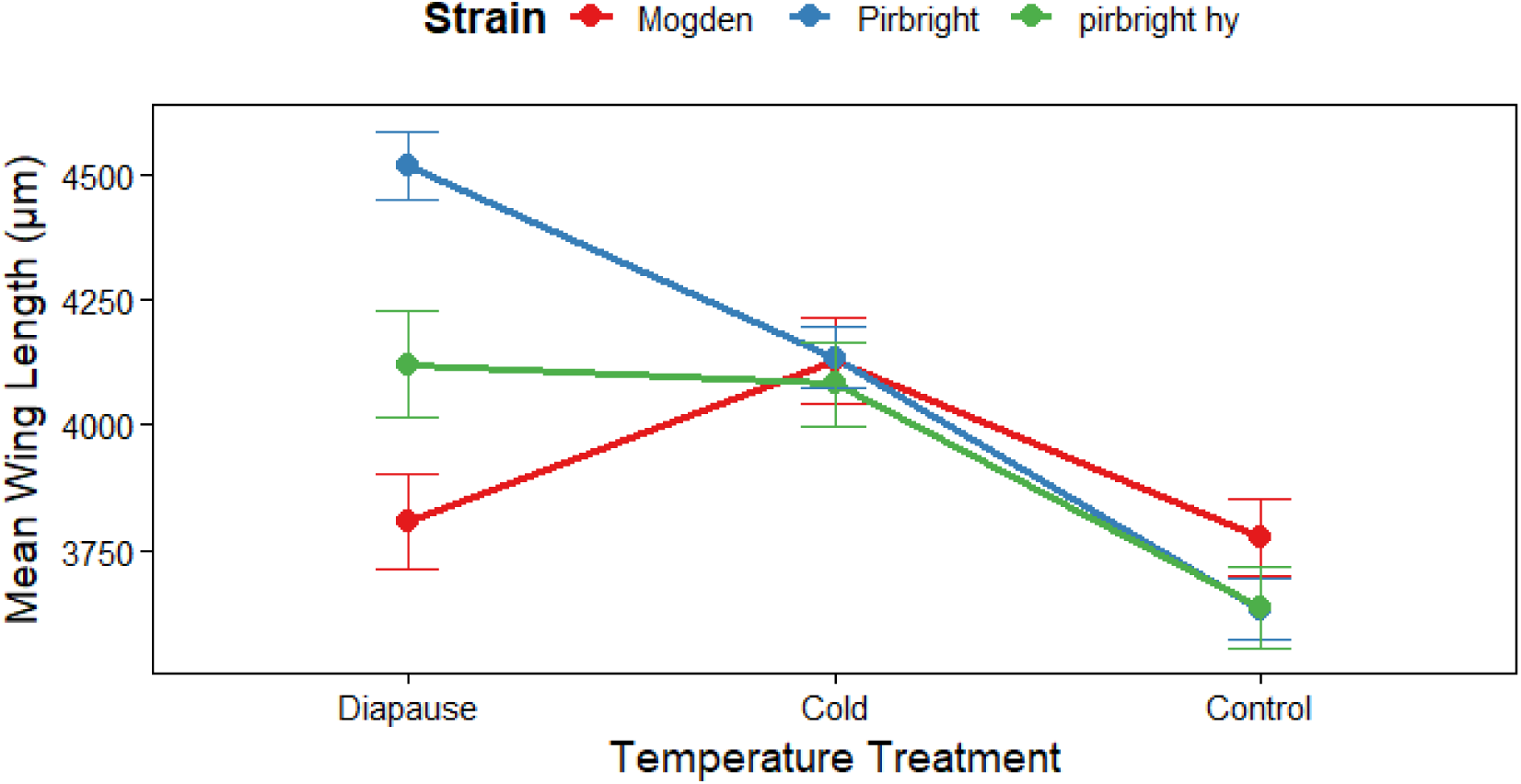
Interaction between temperature and strain on wing size. Treatment × strain interaction plot showing mean ± SE wing length for each strain (Mogden, Pirbright, and Pirbright Hybrid) across diapause, cold, and control temperatures. Non-parallel slopes demonstrate a significant interaction between temperature and genetic background, indicating strain-specific thermal plasticity in wing development

### 3.3 Effect of temperature on spermatheca size

Spermatheca size was strongly influenced by temperature (Figure 9). Females reared under diapause-inducing conditions (10 °C) exhibited markedly reduced spermatheca size compared with those reared at 14 °C and 26–27 °C.

**Figure 9.**
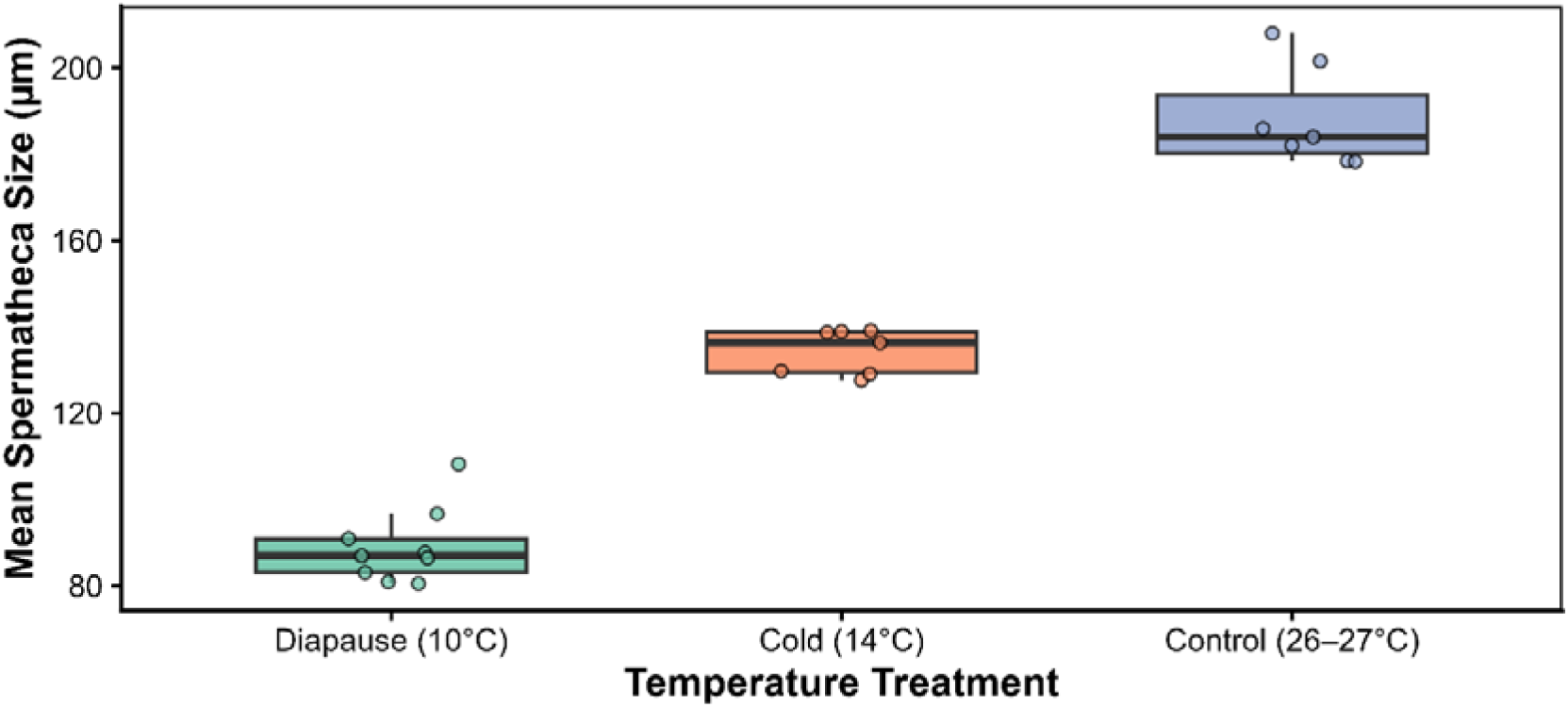
Effect of temperature on spermatheca size in *Culex pipiens*. Mean spermatheca diameter (µm) of females reared at diapause (10 °C), cold (14 °C), or control (26–27 °C) temperatures. Each data point represents one female (mean of three spermathecae), and boxplots summarise the distribution per treatment. Spermatheca development was strongly reduced at 10 °C, intermediate at 14 °C, and fully developed under control temperatures, confirming temperature-induced reproductive arrest.

One-way ANOVA revealed a highly significant effect of temperature on spermatheca size (F□, □□ = 242.4, p = 9.55 × 10□¹□). Post hoc Tukey comparisons confirmed that all pairwise differences between treatments were statistically significant (p < 0.001).

Females reared at 14 °C showed intermediate spermatheca development, whereas those reared under control conditions exhibited fully developed reproductive structures. These results indicate strong inhibition of reproductive development under diapause-inducing conditions.

### Integrated response of wing size and spermatheca development across temperature treatments

When examined together, wing size and spermatheca development exhibited opposing responses across temperature treatments (Figure 10). Wing size increased progressively from control to cold and diapause conditions, whereas spermatheca development decreased across the same gradient. This produced a clear divergence between somatic and reproductive traits under diapause-inducing conditions. A strong negative association (r = −0.95) between wing size and spermatheca development provides direct evidence for a trade-off between somatic growth and reproductive investment.

**Figure 10.**
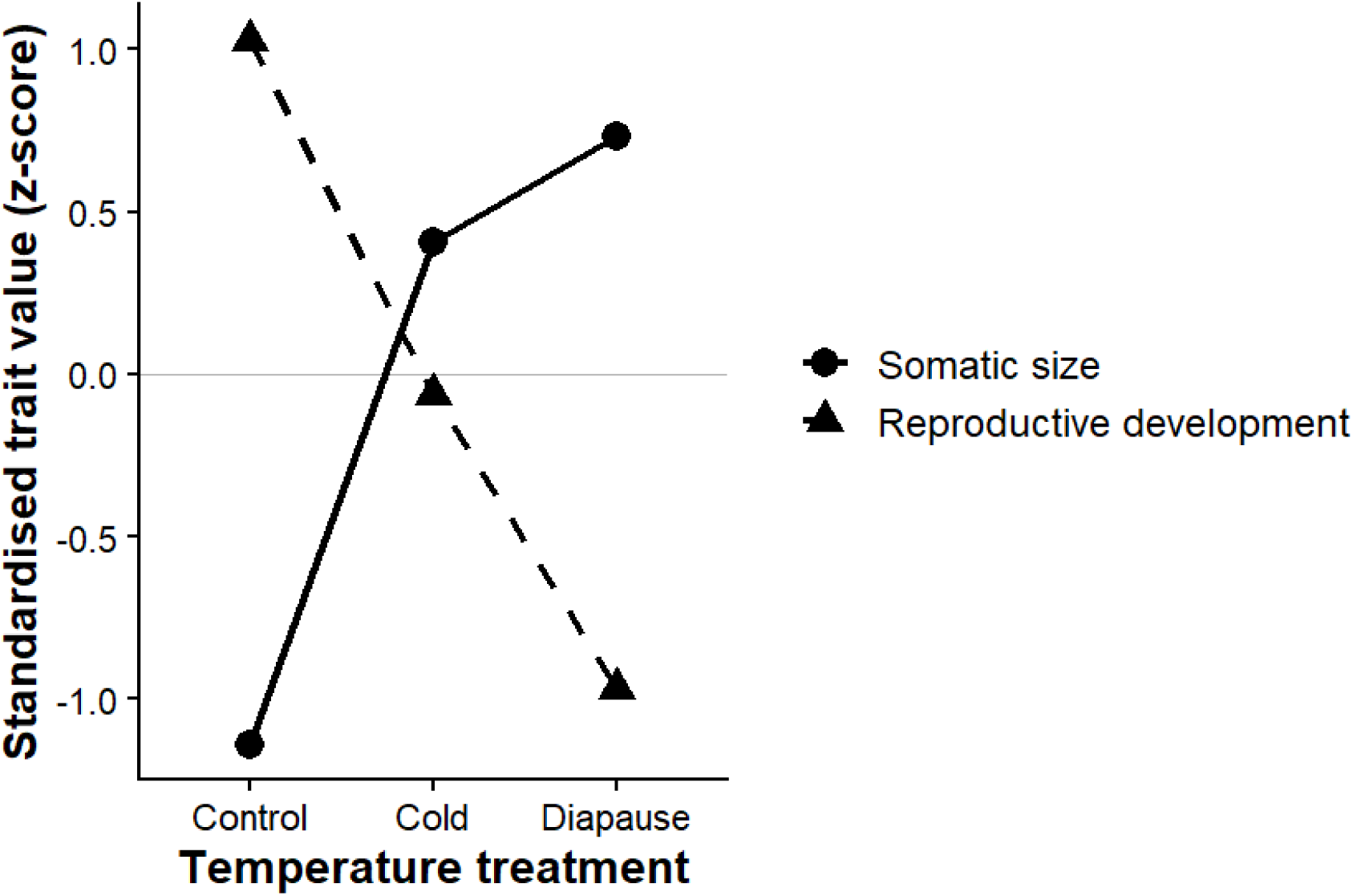
Opposing trajectories of somatic and reproductive traits across temperature treatments in *Culex pipiens*. Standardised mean trait values (z-scores) for wing size and spermatheca development across control, cold, and diapause-inducing conditions. Somatic size increased progressively under colder conditions and was highest in diapause-reared mosquitoes, whereas reproductive development declined across the same gradient and was lowest under diapause-inducing conditions. These opposing responses indicate a coordinated shift from reproductive investment towards somatic maintenance under diapause conditions.

## Discussion

This study investigated the effects of temperature on diapause expression in *Culex pipiens* across behavioural, morphological, and reproductive traits. Overall, diapause-inducing conditions (10 °C) combined with a short photoperiod resulted in strong suppression of blood-feeding, increased somatic size, and marked inhibition of reproductive development. In contrast, mosquitoes reared at 14 °C exhibited intermediate phenotypes, while those maintained under control conditions showed normal feeding and reproductive activity.

Additionally, responses varied across strains: *Culex pipiens pipiens* and the hybrid strain (both overground populations) exhibited similar diapause responses, whereas the Mogden (*Cx. p. molestus*) strain showed a reduced or absent diapause response, indicating clear differences in diapause plasticity between strains. Crucially, the opposing responses of somatic and reproductive traits provide empirical evidence that diapause involves reallocation of resources away from reproduction towards survival.

These findings are particularly important given that *Culex pipiens* is a primary vector of several medically important arboviruses, including West Nile virus and Usutu virus, both of which have been found across Europe (Simonin, 2024). As transmission risk increases, there is a growing need to understand how diapause plasticity within and between strains influences mosquito survival and seasonal transmission dynamics.

Previous studies have shown that diapause expression in *Cx. pipiens* is strongly influenced by genetic background, with different populations and subspecies exhibiting varying sensitivity to diapause-inducing cues (Sauer et al., 2022). However, direct empirical comparisons of multiple strains under controlled environmental conditions remain limited. Understanding such strain-specific responses is essential for predicting overwintering success, the timing of spring emergence, and ultimately the seasonal dynamics of arbovirus transmission.

Blood-feeding inhibition was one of the clearest indicators of diapause, with near-complete suppression observed in females reared at 10 °C across both age groups. The absence of a significant interaction between temperature and age suggests that diapause-associated behavioural arrest is established during development and maintained throughout adult life.

This finding aligns with classical descriptions of diapause in *Culex pipiens*, in which females prioritise energy conservation over host-seeking behaviour (Schäfer and Lundström, 2001; Robich and Denlinger, 2005). The observed suppression of blood-feeding has important implications for arbovirus transmission because blood-feeding is required for horizontal transmission between hosts and mosquitoes, the near-complete inhibition of feeding observed under diapause-inducing conditions would be expected to substantially reduce transmission opportunities during winter.

Morphological responses to temperature were reflected in wing size variation. Mosquitoes reared under diapause-inducing conditions exhibited the largest wings, followed by those reared at 14 °C, while control mosquitoes had the smallest wings. This pattern is consistent with the temperature–size rule, whereby lower developmental temperatures result in increased adult body size due to prolonged developmental periods. Larger body size in diapausing females may confer adaptive advantages, including enhanced energy storage and increased overwinter survival. The significant interaction between temperature and strain further indicates that thermal plasticity is not uniform across populations (Schäfer and Lundström, 2001; A Kassim *et al*., 2013; Nelms *et al*., 2013), with Pirbright and hybrid strains showing strong responses, while the Mogden strain exhibited limited variation. This reduced sensitivity is consistent with previous observations that *Cx. p. molestus* populations do not exhibit a classical diapause response (Nelms et al., 2013).

Reproductive development, assessed via spermatheca size, showed the strongest response to temperature. Females reared at 10 °C displayed severely reduced spermathecae, consistent with reproductive arrest, while those at 14 °C exhibited intermediate development. This supports the interpretation that 14 °C induces a partial or incomplete diapause phenotype, rather than a fully active reproductive state. Together, these findings demonstrate that spermatheca size is a highly sensitive indicator of diapause status.

Previous studies have reported reduced diapause induction in long-established laboratory colonies of *Culex pipiens*, likely due to the loss of diapause responsiveness following prolonged maintenance under stable laboratory conditions without reintroduction of wild genetic variation (Field et al., 2022). In contrast, the colony used in this study is relatively recently established, which may better preserve natural diapause-associated traits. This difference in colony history may explain the stronger and more consistent diapause responses observed here, suggesting that colony age and maintenance practices play a critical role in shaping diapause expression in laboratory populations.

Importantly, the observed strain-specific differences indicate that diapause plasticity varies among commonly used laboratory populations. Such variation is likely to influence overwintering survival, the timing of spring emergence, and the synchronisation of host-seeking behaviour with environmental conditions. These factors are directly relevant to arbovirus transmission dynamics, as diapause determines the persistence of vector populations between transmission seasons.

The opposing trajectories of wing size and spermatheca development across temperature treatments provide evidence for a trade-off between somatic and reproductive investment during diapause. Under diapause-inducing conditions, mosquitoes exhibited increased body size alongside pronounced suppression of reproductive development, consistent with a reallocation of resources away from reproduction and towards survival. This coordinated shift highlights diapause as an integrated life-history strategy, in which physiological and morphological traits are jointly regulated to optimise overwinter persistence and shape seasonal population dynamics under unfavourable environmental conditions.

## Limitations of the study

Several limitations should be considered when interpreting these findings. First, temperature and photoperiod were experimented on simultaneously to generate diapause-inducing conditions. While this approach reflects the natural environmental cues that induce diapause in *Culex pipiens*, it prevents the independent effects of temperature and photoperiod from being separated. Consequently, the observed responses should be interpreted as integrated responses to the combined diapause-inducing environment rather than to temperature alone.

Secondly , the study utilised long-established laboratory strains. Although the relatively strong diapause responses observed here suggest that key diapause-associated traits have been retained, laboratory populations may not fully represent the phenotypic diversity and environmental responsiveness of wild mosquito populations.

In addition, sex-specific responses were not considered in this study. Only female mosquitoes were examined because of their direct relevance to blood-feeding and pathogen transmission. However, male mosquitoes may exhibit different physiological responses to diapause-inducing conditions, and future studies should investigate whether diapause-associated plasticity differs between sexes.

Finally, infection status was not assessed. While this study demonstrates substantial behavioural, morphological, and reproductive changes associated with diapause, it does not directly determine whether arboviruses persist within mosquitoes during diapause or how infection may influence diapause expression.

Future research should address these limitations by experimentally separating temperature and photoperiod effects, incorporating recently collected field populations, and increasing biological replication. Measurements of lipid accumulation, energy reserves, and metabolic activity would further strengthen understanding of the physiological mechanisms underlying diapause. Most importantly, future studies should investigate viral persistence during diapause and the consequences of diapause termination for arbovirus transmission. Determining whether infected mosquitoes can survive winter and subsequently resume transmission in spring is central to understanding the overwintering ecology of viruses such as West Nile virus and Usutu virus.

These future directions provide an important link between diapause physiology and arbovirus transmission. While diapause suppresses blood-feeding and therefore limits opportunities for horizontal transmission, it may simultaneously facilitate long-term pathogen persistence by increasing vector survival. Understanding this balance between transmission suppression and overwinter survival remains a key challenge in predicting arbovirus persistence in temperate regions under environmental change.

By demonstrating coordinated shifts across behavioural, morphological, and reproductive traits, this study moves beyond reductionist approaches that characterise diapause using single traits. Instead, our findings support a framework in which diapause is an integrated life-history strategy governed by resource allocation trade-offs between survival and reproduction. This perspective provides a mechanistic basis for understanding mosquito persistence and offers a foundation for incorporating diapause dynamics into predictive models of arbovirus transmission under environmental change.

## Conclusion

Diapause in *Culex pipiens* is a coordinated, multi-trait response driven by temperature and genetic background. Diapause-inducing conditions resulted in suppressed blood-feeding, increased body size, and reduced reproductive development, while intermediate temperatures produced partial phenotypes, indicating a continuum rather than a binary state. The opposing responses of wing size and spermatheca development highlight a trade-off between somatic growth and reproduction, reflecting a shift towards survival during unfavourable conditions. Variation among strains further demonstrates that diapause plasticity is population dependent. Together, these findings emphasise the importance of considering multiple traits when assessing diapause and its implications for mosquito persistence and seasonal disease risk.

